# A Novel Angiotensin-Converting Enzyme 2 Fusion Protein Reduces Cardiotoxic Effects of Doxorubicin

**DOI:** 10.64898/2026.11.26.769584

**Authors:** Damian N. Di Florio, Naveen K. H. Nagaiah, Capella R. Weems, Isaac E. Prah, Olivia A. Sloan, Christopher J. Ray, Pooja P. Advani, Matthew R. Fleming, Philip A. Romero, Pete Heinzelman, Nadine Norton

**Author notes:** Corresponding author. Name: Nadine Norton, 4500 San Pablo Road, Jacksonville, FL, 32224. These authors contributed equally to this work (Damian N. Di Florio, Naveen K. H. Nagaiah). Senior Authors (Matthew R. Fleming, Philip A. Romero, Pete Heinzelman, Nadine Norton).

## Abstract

**Background:** Anthracyclines, such as doxorubicin, are a cornerstone of chemotherapy and are the first-line therapy for high-risk breast cancer. Doxorubicin, however, induces dose-dependent cardiotoxicity that can lead to acute and chronic heart failure. In clinical studies with serial measurement of left ventricular ejection fraction (LVEF), targeting the Renin-Angiotensin System (RAS) with angiotensin-converting enzyme (ACE) inhibitors was effective in ≈ 40% of patients (responders), but in 45% of patients, there was no response in LVEF. As such, there is a critical need to find new cardioprotective strategies. We posited that efficacy is limited by sole inhibition of the Angiotensin-II Type 1 Receptor (AT1R) RAS arm without activation of the mas receptor (MasR) RAS arm.

**Methods:** To address this issue, we used a novel engineered angiotensin-converting enzyme 2 (ACE2) fusion protein therapeutic, which possesses enhanced activity and substrate specificity compared to wild-type ACE2 and contains a human IgG Fc domain that increase**s** circulation lifetime; this approach simultaneously attenuates AT1R activation and activates MasR.

**Results:** Mice treated with this protein, known as ACE2dm-Fc, in conjunction with doxorubicin administration had increased longitudinal body weight gain and were protected from left ventricular dysfunction and cardiac remodeling compared to mice receiving doxorubicin alone. Additionally, bulk-tissue RNA sequencing identified gene expression correlates of heart function that distinguished mice receiving both ACE2dm-Fc and doxorubicin from those given only doxorubicin.

**Conclusions:** Our results establish ACE2dm-Fc’s potential as a novel treatment for doxorubicin-induced decline in LVEF.

**Clinical Perspective:**

- Attempts to modulate renin-angiotensin signaling (RAS) to provide cardioprotection against doxorubicin induced cardiotoxicity have yielded mixed results in the clinic.
- Our approach to RAS modulation is mechanistically unique to prior therapeutics and has not been assessed in the context of doxorubicin induced cardiac dysfunction.
- We demonstrated that early treatment with this novel fusion protein therapy provided functional protection against doxorubicin induced heart injury, demonstrating pre-clinical promise for future prophylactic use in cancer patients treated with doxorubicin to reduce the rate and severity of cardiac dysfunction.

## INTRODUCTION

Anthracyclines are incorporated into > 50% of chemotherapy regimens and remain first-line therapy for several adult and pediatric cancers, including leukemia, sarcoma, lymphoma, and high-risk breast cancer. ^1,2^ Among them, doxorubicin is highly effective for breast cancer^3,4^ and is one of the few effective options for treating metastatic triple-negative metastatic breast cancer, ^5^ an aggressive disease with a five-year survival rate of only approximately 10 percent.^6^ Doxorubicin’s clinical utility is limited by dose-dependent cardiotoxicity that can manifest as dilated cardiomyopathy, arrhythmia, and heart failure (HF).^7,8^ The incidence of doxorubicin-induced heart failure rises sharply with cumulative dose (3% < 400 mg/m² to 18% with ≥550 mg/m²).^9,10^ Further risk is conferred by co-administration of the anti-HER2 monoclonal antibody drug trastuzumab, ^11^ patient cardiovascular risk factors, ^12–15^ and subclinical cardiac dysfunction.^16^ Dexrazoxane is the only FDA-approved drug for mitigating doxorubicin cardiotoxicity^17,18^ but its use is limited to patients with metastatic breast cancer with ≥ 300 mg/m^2^ exposure profiles.^19^ Despite this efficacy, dexrazoxane is rarely administered in adult practice: in a national cohort of 13,331 adults receiving frontline anthracycline therapy for hematologic malignancy, dexrazoxane was given to 0.37% of patients with diffuse large B-cell lymphoma and 0.14% of those with de novo acute myeloid leukemia, with use concentrated in a small number of practices.^20^

Although standard-of-care HF treatments have been pursued as agents to ameliorate doxorubicin cardiotoxicity, clinical study outcomes indicate that these drugs do not adequately attenuate doxorubicin’s cardiotoxic effects. Beta-blockers, intended to provide cardiac benefits by preventing interaction between adrenalins and their receptors on cardiac cells, and often used as anti-hypertensives for HF patients, had no effect on all cause morbidity, cardiac function and quality of life when given to cancer patients receiving doxorubicin chemotherapy.^21^ In another trial, the angiotensin converting enzyme (ACE) inhibitor enalapril, which has anti-hypertensive properties that could preserve cardiac function in doxorubicin patients by reducing levels of the pro-inflammatory, blood pressure-elevating Renin-Angiotensin system (RAS)**-**associated angiotensin-II (Ang-II) peptide in blood and/or cardiac tissue, did not protect breast cancer patients from doxorubicin cardiotoxicity.^22^ Similarly reflective of existing HF drugs’ limitations in preventing doxorubicin cardiotoxicity, breast cancer patients treated with doxorubicin and given the Ang-II type1 receptor (AT1R) antagonist candesartan, which is often prescribed for hypertension management, did not show improved cardiac function relative to control subjects in combination with beta-blockers.^23^ Most recently, the PRADA II trial tested sacubitril/valsartan, an angiotensin receptor-neprilysin inhibitor combination that augments natriuretic peptide signaling while blocking AT1R, in 138 women receiving anthracycline-based adjuvant breast cancer therapy; despite this dual mechanism, the decline in left ventricular ejection fraction was not significantly attenuated relative to placebo, although measures of myocardial deformation and injury improved.^24^ In practice, the blood pressure-lowering effects intrinsic to these agents often limit their initiation and uptitration in patients receiving cardiotoxic chemotherapy, who are frequently hypotensive from concurrent treatment; in troponin-positive patients receiving anthracyclines, sacubitril/valsartan was associated with symptomatic hypotension in 14% versus 1.8% with placebo.^25^

Both ACE inhibitors and AT1R antagonists modulate the RAS by reducing AT1R activation and thus attenuating the vasoconstriction, inflammation, fibrosis, and oxidative stress that such activation can promote. This mechanism of action overlooks the potential to beneficially modulate the RAS by increasing activation of the mas receptor (MasR), which is vasodilative, anti-inflammatory, and anti-oxidative,^22,26–30^ via increasing levels of this receptor’s angiotensin-(1-7) Ang-(1-7) peptide cognate agonist. We sought to use angiotensin-converting enzyme 2 (ACE2), a carboxypeptidase that removes the C-terminal amino acid of the Ang-II octapeptide to generate the Ang-(1-7) septapeptide, as an agent that simultaneously activates MasR and reduces AT1R activation. In particular, we employed an engineered human ACE2 T371L/Y510I double mutant,^31^ referred to as ACE2dm, that possesses increased specificity toward Ang-II relative to the wild-type ACE2 enzyme. This specificity can reduce ACE2’s propensity to cause side effects due to hydrolysis of the numerous off-target peptides, such as apelin peptides that are important for cardiac function, that are hydrolyzed by wild type ACE2. To further enhance the clinical relevance of this candidate therapeutic, a human IgG1 fragment crystallizable (Fc) domain was genetically fused to the ACE2dm enzyme domain to increase circulation lifetime as reported in mouse studies with non-dm recombinant ACE2s.^32^ The resulting ACE2dm-Fc fusion protein was then evaluated in a mouse model of doxorubicin-induced cardiotoxicity as an initial step to assess its merits as a cardioprotective agent targeting doxorubicin’s negative cardiac effects in cancer patients, particularly those with high-risk breast cancer.

## METHODS

### Study Design

We chose n = 10 mice per group for this study based on previous work showing that this group size allowed differences in heart function to be observed.^33,34^ For RNAseq, n = 6 per group was used and samples were chosen on the basis of proximity to the mean LVEF % such that they were phenotypically representative of their respective groups. We did not exclude data or apply outlier exclusion. A single endpoint was chosen for the study and we did not repeat experiments. Animal subjects (female, 8-week-old FVB/NJ) were used in this study. The primary objective of the study was to determine if ACE2dm-Fc treatment provided resistance against systolic dysfunction following doxorubicin administration. We used a vehicle group (receiving PBS vehicle injections for doxorubicin and ACE2dm-Fc) as a stress-matched control, doxorubicin only group as a functional pathological group, and a group receiving doxorubicin and ACE2dm-Fc treatment concurrently. Mice (Jackson Laboratories) were randomized to one of the three groups. One laboratory personnel performed echocardiography and another, blinded to sample IDs, performed analysis (i.e. tracing echo images). No other assays or evaluations were performed using blinding. Additional details for methodologies and materials are detailed below and in the Supplemental Methods.

### Generation of ACE2dm-Fc

The ACE2dm mutations T371L/Y510I were introduced into a vector encoding the wild type ACE2-Fc fusion protein (Addgene ID 145163),^35^ and used for transient transfection of Expi293 cells (LifeTech). Affinity purification was performed using mAb Select (IgG Fc-binding) resin prior to size exclusion chromatography purification using a Cytiva Sephacryl S-300 exclusion column. SEC column eluate was concentrated with a 100 kDa MWCO Amicon centrifugal concentrator. The final product was stored in PBS with 3uM ZnCl_2_ added to preserve enzyme activity. A more detailed description of this procedure and protein quality control (SDS-PAGE and SEC) can be found in the Supplemental Methods.

### Mouse Model

Female FVB/NJ WT Mice (Jackson Laboratories) at 8 weeks of age were given doxorubicin (4 mg/kg) on days 0, 5, 10, 13, 16, 19 concurrent with ACE2dm-Fc treatment (3 mg/kg) on days 0, 5, and 10. Both doxorubicin and ACE2dm-Fc were administered by intraperitoneal injection. Treatment groups consisted of mice receiving PBS vehicle in place of both doxorubicin and ACE2dm-Fc (VEH), mice receiving doxorubicin and PBS vehicle in place of ACE2dm-Fc (DOX), and mice receiving doxorubicin and ACE2dm-Fc (DOX + ACE2dm-Fc). Bodyweight measurements were taken 1-2 times weekly and systolic heart function was assessed using echocardiography at day 55. Mice were euthanized the following day. In brief, mice were anesthetized with 80:10 mg/kg ketamine:xylazine with an induction period of 7 minutes per IACUC guidelines.

### Echocardiography

Systolic heart function was assessed by short-axis motion-mode echocardiography using a Vevo 3100 Preclinical Ultrasound System equipped with an MX550D transducer. Mice were anesthetized with isoflurane prior to transfer to a Vevo Imaging Station platform. Mouse hair was removed using Nair. Parasternal short-axis views were acquired at the mid-left ventricle at the region of greatest radial papillary extension into the left ventricle. Motion(M)-mode traces were analyzed in Vevo Labs (version 5.9.0) using three cardiac cycles between breaths. Representative M-mode traces were time-adjusted to show three cardiac cycles. Additional parameters not shown in the main text figures appear in Fig. S1.

### Histology and Myocardial Fibrosis

After excising mouse hearts, sections were fixed in 4% PFA for 24 hours prior to replacement with PBS solution and storage at 4 C. Tissue was embedded in paraffin blocks and cut at a thickness of 5 µm. Slides were stained using Masson’s Trichrome to evaluate ventricular myocardial fibrosis. Slides were scanned at 40x using an Aperio AT2 Slide Scanner (Leica). Aperio Image Scope’s Positive Pixel Count Algorithm was used to determine staining intensity indicative of collagen deposition in ventricular tissue. Mean positive pixel intensity (MPI) was calculated using the algorithm’s “Iavg” output, which represents a raw light intensity (i.e. lower values indicate greater strength of staining as less light passes through them); the Iavg value was subtracted from the maximum intensity value (255) to determine MPI (MPI = 255-Iavg).

### Gene Expression

Heart tissue was snap frozen in liquid nitrogen and later thawed for RNA extraction. Tissue homogenates were prepared using Dounce homogenization at 200 mg/mL in tris-buffered saline with protease and phosphatase inhibitors (cOmplete and PhoSTOP, Sigma-Aldrich). A volume of 40 µL of the resulting homogenate was lysed with 350 µL buffer RLT (supplemented with β-mercaptoethanol) prior to genomic DNA removal using gDNA eliminator columns (Qiagen). The flow through was processed with a RNeasy Mini Kit (Qiagen) per the manufacturer’s instructions. RNA was eluted in nuclease free water and quantified using a NanoDrop spectrophotometer.

Subsequently, 2 µg of RNA was used to generate cDNA using a high-capacity cDNA reverse transcription kit (ThermoFisher). Quantitative PCR was performed using Taqman probes (ThermoFisher) and a Quant Studio 3 qPCR device (Applied Biosystems). Expression of *nppa* (Mm01255747_g1), *nppb* (Mm01255770_g1), *myh6* (Mm00440359_m1), *myh7* (Mm00600555_m1), *col1a1* (Mm00801666_g1), *col3a1* (Mm00802300_m1), *polr2a* (Mm00839502_m1), *cd38* (Mm00483143_m1), *acta1* (Mm00808218_g1), *hmox1* (Mm00516005_m1), *sod2* (Mm01313000_m1), and *gpx4* (Mm04411498_m1) was quantified with the *polr2a* gene being used for normalization. Expression fold changes were determined relative to the VEH mouse group.

### Western Blots

200 µL of heart tissue homogenate was combined with 50 µL 5x RIPA buffer and incubated on ice for 30 minutes prior centrifugation at 21,000xg at 4 C. Supernatant was collected and protein concentration determination using a BCA Assay (Pierce). For blots, 30 µg of lysate was combined with Laemmli buffer (diluted to 1X) and denatured at 95 C for 5 min. Denatured lysate was loaded onto 4-12% Bis-Tris gradient gels (ThermoFisher) and run at 200V for 35 minutes in MES buffer (Invitrogen). Protein was transferred to PVDF membranes using an iBlot3 (Invitrogen) prior to blocking with 1x iBind Flex solution (Invitrogen). Primary and secondary HRP-conjugated antibody incubations were performed using the iBind Flex Western System (Invitrogen). The following primary antibodies were used: Vinculin (Sigma, V9131, 1:4,000), Angiotensin-(1-7)-Mas Receptor (ThermoFisher, AAR-013, 1:200), and Phospholamban (ThermoFisher, MA3-922, 1:4,000). Anti-mouse and anti-rabbit secondary antibodies (Jackson Labs) were used at 1:8,000. The SuperSignal West Pico PLUS Chemiluminescent Substrate (ThermoFisher) was used along with a ChemiDoc Imaging System (BioRad) to image blots. Blots were assessed with densitometry analysis (ImageJ) relative to vinculin loading control and normalized to the VEH group. Representative blot images appear in the main figures, and all samples and uncropped blots can be found in Fig. S2.

### Bulk Tissue RNA Sequencing and Analysis

Sample RIN scores were determined using an Agilent 2100 Bioanalyzer with RNA Nano 6000 kits (Agilent) according to manufacturer instructions. Six biological replicates per group were used for RNA sequencing with selection on the bases of QC (RIN > 7, 260/280 > 1.8 and 260/230 > 1.5) and proximity to mean LVEF % for each group. Sequencing and initial analysis to generate gene expression matrices was performed by PlasmidSaurus. Gene expression matrices were used to perform differential expression analysis of protein coding genes using DESeq2.^36^ sPLS-DA was performed using mixOmics.^37,38^ Various gene groups (from Boolean / overlap analysis) were used to perform multiple linear regressions and correlation data in the manuscript is displayed at standardized (std.) beta, nominal P, and Benjamin-Hochberg False Discovery Rate (FDR). Data was re-plotted in GraphPad Prism. A detailed description of RNAseq QC information for samples, sample selection, and the PlasmidSaurus pipeline can be found in the Supplemental Methods.

### Statistics

For survival analysis, a log-rank, i.e., mantel-cox, test was used. To determine change in body weight over time, percent change from baseline body weight (Δ%) was used and a mixed-effects model (REML; fixed effects: time, treatment, treatment x time) was applied. Holm-Sidak multiple comparisons were used to compare groups each day of measurement. Linear regression was performed pairwise to determine differences in slopes between groups. When slopes were not significantly different, follow up testing for differences in y-intercepts was performed. For all other data (besides RNAseq), groups were compared using a 1-way ANOVA with Holm-Sidak corrected pairwise tests. Data with error bars are represented with mean ± SEM. Statistical methods used for RNA sequencing are detailed in the Methods section for ‘Bulk Tissue RNA Sequencing and Analysis’ and defined throughout the manuscript.

### Independent Data Access and Analysis

DND had full access to all the data in the study and takes responsibility for its integrity and the data analysis.

### Materials and Data Availability

Data used for RNA sequencing analysis is supplied as a gene expression matrix and DeSeq2 outputs (for each comparison) and all other data used for the generation of figures in this study are available in the Supplemental Data. No new code was generated for this study. Fastq files will be uploaded to GEO following initial review and made publicly available upon publication of the manuscript. Any other information or supporting data not included is available upon reasonable request.

## RESULTS

### Study Timeline and Weight Changes in Response to Doxorubicin Treatment

To ensure translational relevance with respect to sex for breast cancer patients treated with doxorubicin,^39–41^ we used 8-week-old female FVB/NJ mice in our study. Mice were given doxorubicin (or PBS vehicle control) via intraperitoneal injection (4 mg/kg) on study days 0, 5, 10, 13, 16, and 19 (cumulative dose = 24 mg/kg). A third group of animals was given doxorubicin treatment concurrent with intraperitoneal injection of ACE2dm-Fc (3 mg/kg) on days 0, 5, and 10 (**Fig. 1A**). As previous studies found that the blood half-lives of non-dm ACE2-Fc fusion proteins were approximately seven days, the dosing schedule was expected to give rise to appreciable ACE2dm-Fc bloodstream concentrations throughout the doxorubicin administration interval.^32^ Among the mice treated with doxorubicin alone (DOX), two expired before the day 56 study endpoint. All vehicle mice (VEH) and mice receiving both ACE2dm-Fc and doxorubicin (ACE2dm-Fc + DOX) survived throughout the course of study (**Fig. 1B**). Although the loss of two mice from the DOX group suggested an ACE2dm-Fc-conferred survival benefit, the difference in survival was not large enough to be statistically significant.

**Fig. 1.**
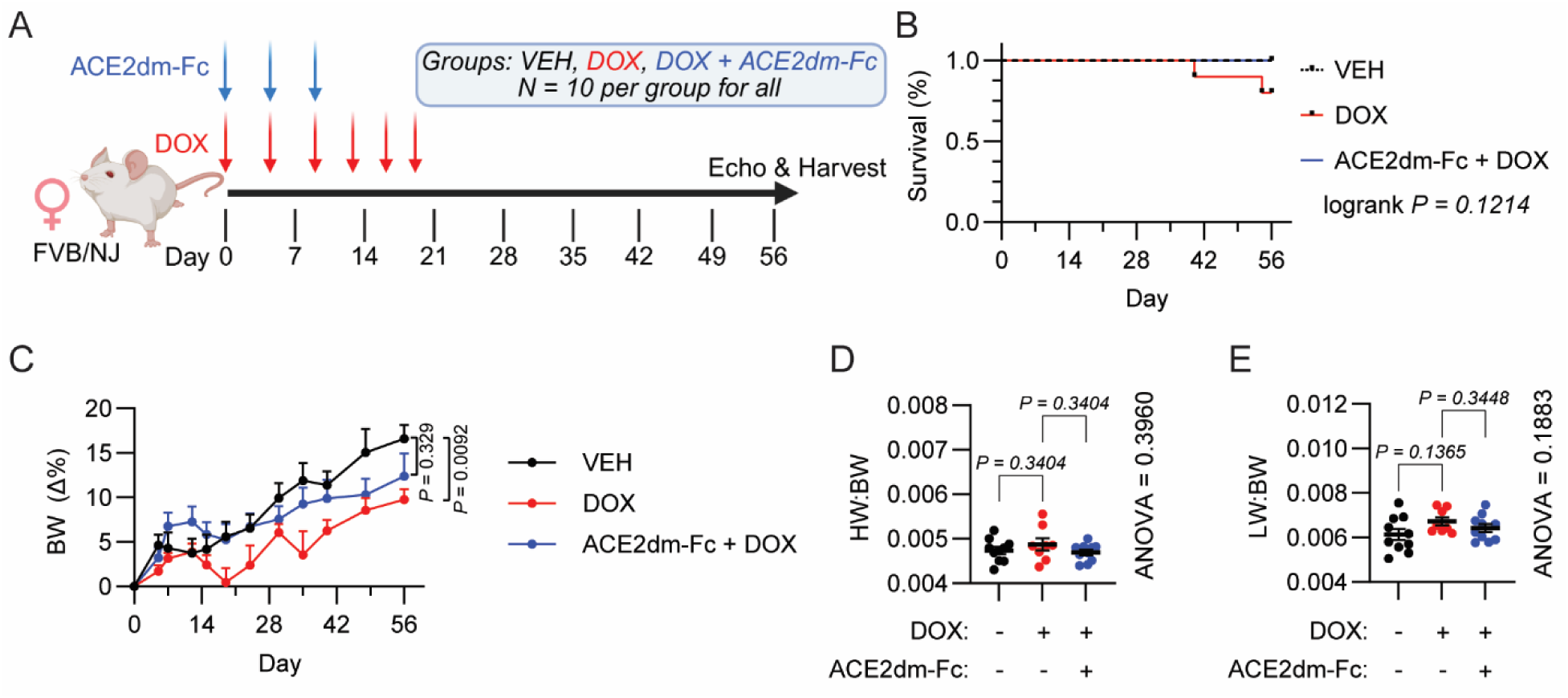
Study design and gross anatomical features of mice. (**A**) Study timeline and dosing schedule (**B**) Survival curve (**C**) Change in animal bodyweight throughout the study period (Δ% denotes change in bodyweight from baseline measurement) (**D**) heart weight to bodyweight (HW:BW) and (**E**) lung weight to bodyweight ratios. Log-rank (mantel-cox) for (**B**), mixed-effects (REML) model for daily comparisons of body weight and linear regression analysis for (**C**), 1-way ANOVA with Holm-Sidak (adjusted p-values shown) follow-up testing comparing VEH vs. DOX and DOX vs. ACE2dm-Fc+DOX in (**D & E**)

At study endpoint, there was significantly less bodyweight increase (Δ%) for DOX than VEH mice (**Fig. 1C**). Bodyweight changes for ACE2dm-Fc + DOX mice (**Fig. 1C**) relative to VEH mice were not significant, thus showing that ACE2dm-Fc promotes the ability to gain bodyweight both during and after doxorubicin administration. As observed in other female mouse models of doxorubicin exposure,^33,42–44^ heart weight and lung weight changes, normalized to bodyweight, were unchanged across groups (**Fig. 1D-E**).

### Assessment of Cardiac Function After Doxorubicin and ACE2dm-Fc Treatments

Both humans and mice experience compromised heart function following treatment with doxorubicin.^41^ To determine whether ACE2dm-Fc treatment conferred functional cardioprotection, we evaluated left ventricular systolic function using short-axis motion (M-mode) echocardiography. ACE2dm-Fc provided resistance to systolic dysfunction imparted by doxorubicin by measures of left-ventricular ejection fraction, fractional shortening and stroke volume (**Fig. 2A-C**), despite lower heart rates for ACE2dm-Fc treated mice **(Fig. S1A)**. Cardiac output was decreased for both the DOX and DOX + ACE2dm-Fc groups **(Fig. S1B)**.

**Fig. 2.**
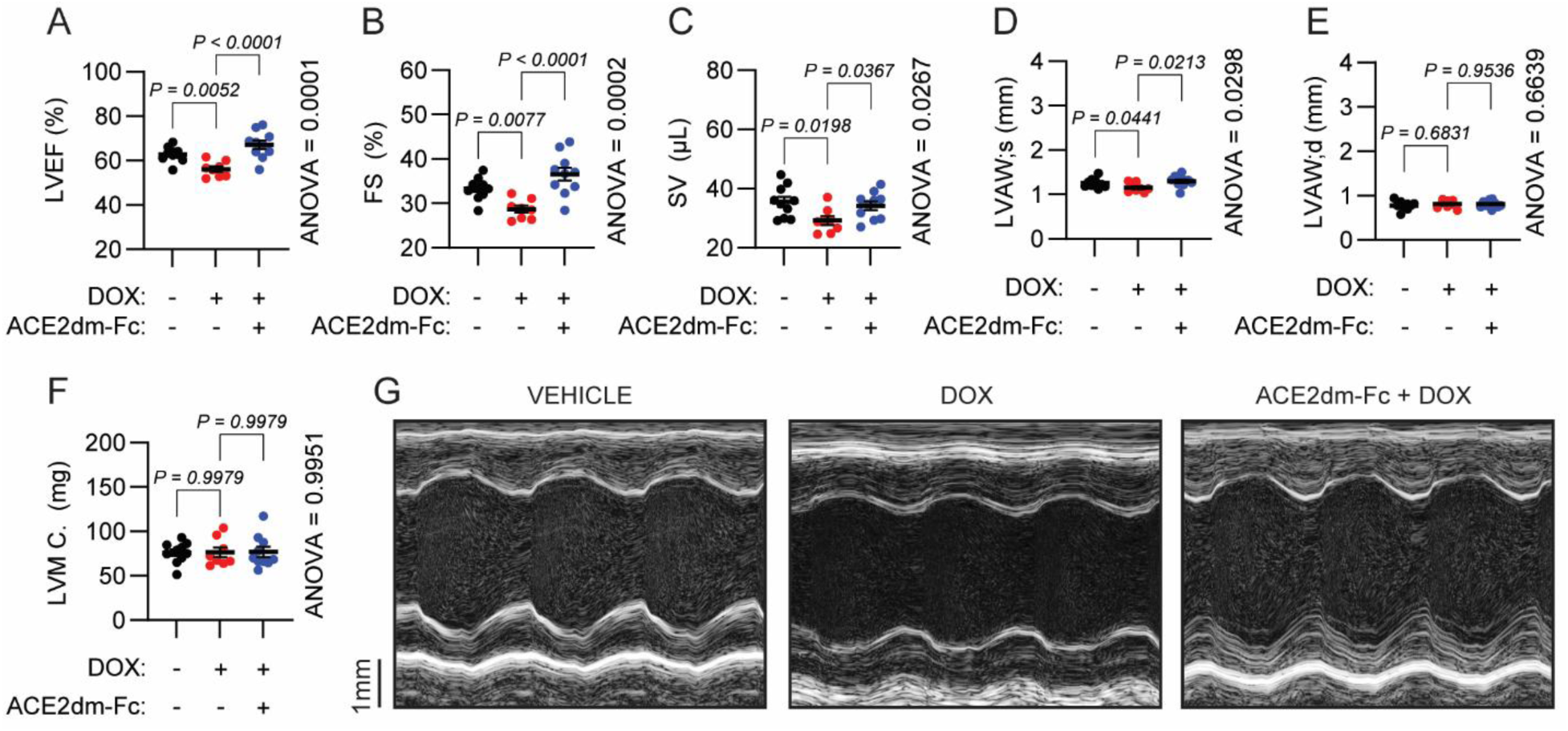
ACE2dm-Fc treatment protects against systolic heart dysfunction induced by doxorubicin. (**A**) Left-ventricular ejection fraction (LVEF), (**B**) fractional shortening (FS), (**C**) stroke volume (SV), (**D**) left-ventricular anterior wall thickness at systole, (**E**) left-ventricular anterior wall thickness at diastole, (**F**) left-ventricular mass corrected (LVM C.) and (**G**) representative M-mode micrographs for study groups time-adjusted to show 3-cycles between breaths. Statistics for all are 1-way ANOVA with Holm-Sidak follow up testing comparing VEH vs. DOX and DOX vs. ACE2dm-Fc+DOX, adjusted p-values are reported.

Left-ventricular anterior wall thinning during systole, which can be reflective of cardiac damage, was observed in DOX mice but not in the ACE2dm-Fc + DOX group (**Fig. 2D**). Wall thickness changes in response to doxorubicin were not observed anteriorly during diastole (**Fig. 2E**), and left-ventricular posterior wall thickness was also unchanged (**Figs. S1C & D**). In accord with the absence of changes in heart weight to bodyweight ratio (**Fig. 1D**), we did not detect differences in estimated left-ventricular mass as determined by echocardiography (**Fig. 2F**). This suggested that the observed prevention of anterior wall thinning by ACE2dm-Fc at systole (**Fig. 2D**) was driven by a difference in radial motion during systole (**Fig. 2A-C**). This observation is congruent with reductions in end-systolic diameter and volume for ACE2dm-Fc treated mice, without differences at end diastole (**Fig. S1E-H**). Echocardiography findings are visually corroborated with representative M-mode images in **Figure 2G**.

### ACE2dm-Fc Reduces Myocardial Fibrosis Following Doxorubicin Administration

To assess whether ACE2dm-Fc + DOX mice were resistant to established gene expression patterns associated with murine doxorubicin cardiotoxicity, we used quantitative PCR to measure transcript levels for genes related to cardiac damage, stress/remodeling, fibrosis, oxidative stress, and senescence. ^45,46^ We observed increased *myh7* expression in response to DOX, which drove down *myh6/7* ratios and was not different with ACE2dm-Fc administration (**Figure 3A**). Similarly, ACE2dm-Fc did not change MasR or phospholamban, which is a marker of impaired cardiac function, levels as determined by Western blot (**Fig. 3B**, fig. S2).

**Fig. 3.**
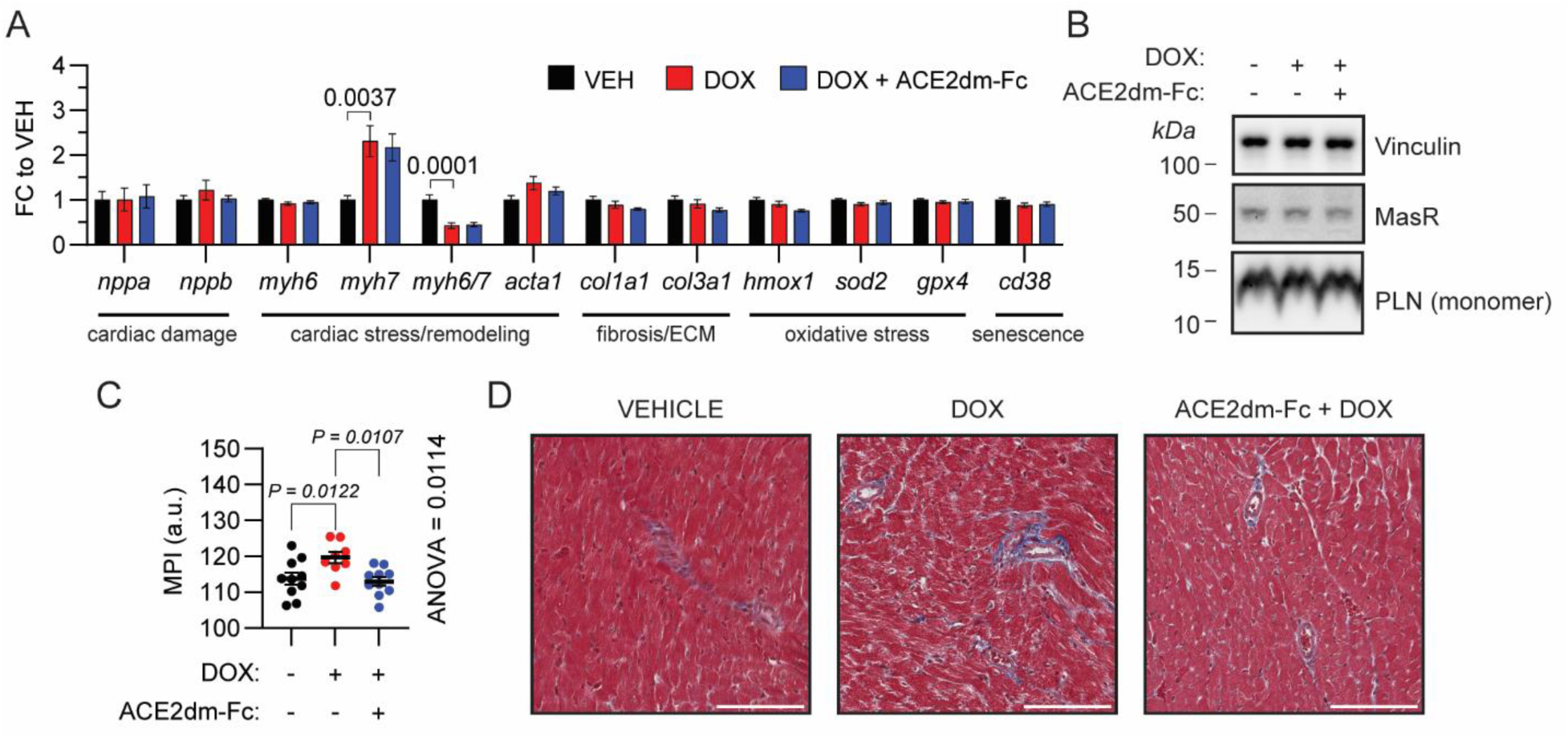
ACE2dm-Fc treatment protects against myocardial remodeling induced by doxorubicin. (**A**) quantitative PCR for various markers (**B**) representative Western blot depicting vinculin (loading control), Mas Receptor (MasR), and phospholamban monomer (PLN). (**C**) Histological analysis of trichrome stained mouse heart sections comparing the mean positive pixel intensity (MPI) and (**D**) representative micrographs (scale bars = 100 µm). Statistics for all are 1-way ANOVA with Holm-Sidak follow up testing comparing VEH vs. DOX and DOX vs. ACE2dm-Fc+DOX, adjusted p-values are reported. Non-significant p-values are not displayed.

In addition to changes in the expression of cardiac stress-associated genes, myocardial remodeling is another major contributor to changes in cardiac function due to doxorubicin exposure. We found that DOX mice had increased fibrosis compared to VEH (**Fig. 3C**). ACE2dm-Fc treatment markedly reduced fibrosis in DOX-treated mice (**Fig. 3C**). These data suggest that the systolic contractile differences across groups in **Fig. 2** may be explained by differences in myocardial flexibility.

### Transcripts Associated with Doxorubicin Cardiotoxicity are Affected by ACE2dm-Fc

To further identify genes with expression affected by ACE2dm-Fc, we performed bulk tissue RNA sequencing (RNA-seq) on hearts collected at the study endpoint. Sparse partial least squares discriminant analysis (sPLS-DA)^37,38^ was carried out using the top 500 most variable protein-coding genes (**Fig. 4A-B**); this analysis enables observation of intragroup clustering that can lead to identification of genes that are differentially expressed across study groups. Component 1 showed separation of mice treated with doxorubicin, and separation for component 2 mouse groups paralleled changes in cardiac function as observed by EC (**Fig. 4B**).

**Fig. 4.**
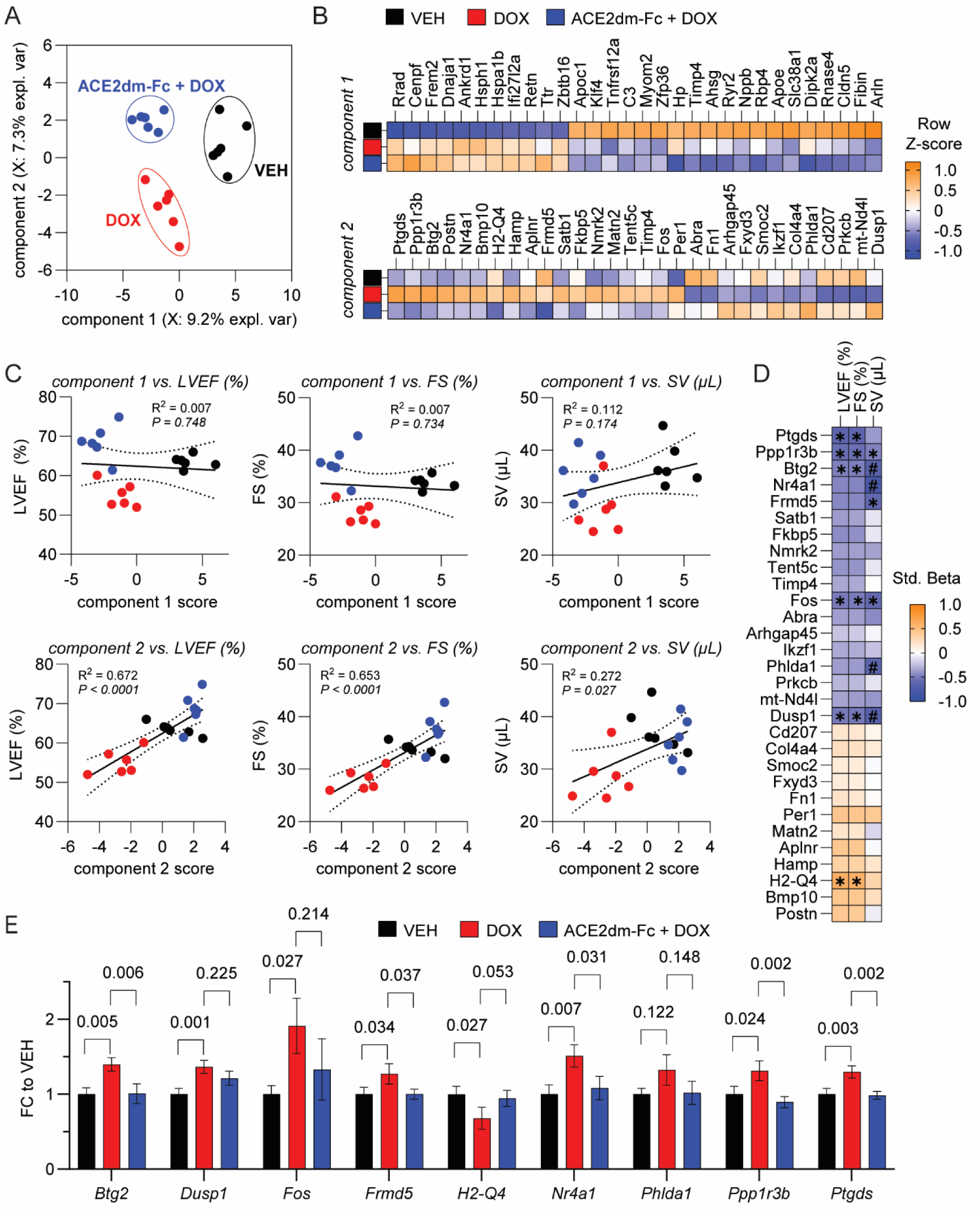
Discriminant analysis identifies transcriptomic correlates of mouse heart function. (**A**) sPLS-DA plot showing separation between groups based on (**B**) genes in components 1 and 2. (LOO-CV balanced error =22.2%, permutation P = 0.005, N = 200 shuffles) (**C**) sPLS-DA component 1 and 2 correlations vs. mouse heart function. (**D**) Component 2 gene correlations vs. mouse heart function. (**E**) Component 2 genes with at least 1 nominal correlation to mouse heart function. For (**C&E**) nominal p values are reported and for component score vs. heart function correlations (simple linear correlation) and pairwise testing from DESeq2 output (Wald test). For (**D**), * signifies a nominal p < 0.05 and # FDR < 0.05 correlation (column adjusted). In component 1 (B, top), for the purpose of figure display, we used the symbol Arln for 1810037I17Rik.

To determine whether component 2 was related to cardiac function, we used component scores to perform correlation analyses versus mouse systolic function (**Fig. 4C**). There were no significant correlations with component 1 scores and heart function and separation along the component 1 axis (**Fig. 4A**) were similar to the directionality of group-level z-scores displayed for component 1 (**Fig. 4B**, top). In contrast, component 2 scores demonstrated significant correlations among phenotypes (**Fig. 4C**, bottom), matching the group-level z-score directionality in the component 2 heat map (**Fig. 4B**, bottom).

To identify individual transcripts associated with changes in echocardiography parameters, we interrogated component 2 gene relationships to cardiac function with linear regression. This analysis revealed 9 genes that were significantly correlated with mouse phenotypes. Four of these genes (*nr4a1*, *phlda1*, *btg2*, and *dusp1*) correlated after phenotype-level false discovery rate (FDR) correction for stroke volume (**Fig. 4D**). No such correlations were observed between transcript level and fractional shortening or ejection fraction. Among the 9 genes with at least nominally significant correlations to systolic function, four of them: *nr4a1*, *btg2*, *ptgds*, *frmd5*, and *ppp1r3b,* exhibited both significantly increased expression when comparing DOX to VEH and significantly decreased expression when comparing ACE2dm-Fc + DOX to DOX (**Fig. 4E**). Based on their inclusion in this set of genes and significant correlation with stroke volume after FDR correction, we identified *nr4a1* and *btg2* as correlates of mouse systolic function for which ACE2dm-Fc treatment provided resistance to transcript changes associated with doxorubicin-induced cardiac dysfunction.

### ACE2dm-Fc Modifies Cardiac Transcript Expression Correlates of Systolic Heart Function

Given that sPLS-DA gene identification was agnostic to fold-change (FC) and/or P-value cutoffs and that *nr4a1* and *btg2* only correlated with stroke volume (rather than multiple systolic function measures), we sought to identify other divergent transcriptomic changes that could correlate with multiple systolic functional parameters. We identified 4 transcript expression differences between VEH and DOX after FDR correction: *lbh*, *rrad*, *nudt4*, and *cenpf* (**Table 1**). When comparing ACE2dm-FC + DOX to VEH, the same 4 transcripts were identified, with changes in the same direction relative to VEH, and 4 additional genes were found to be unique to ACE2dm-Fc + DOX relative to VEH: *frem2*, *zcrb1*, *ifi27l2a*, and *rhou*. Comparing ACE2dm-Fc + DOX to DOX however, did not reveal genes meeting FDR significance. Although *frem2* and *ifi27l2a* showed nominal significance for ACE2dm-Fc relative DOX, their directionality with respect to VEH remained the same for DOX and ACE2dm-Fc + DOX. Collectively, the above results show that, despite the clear impact of ACE2dm-Fc on cardiac function as observed at the study endpoint, the magnitude of ACE2dm-Fc-driven changes in gene transcript levels at this same timepoint was limited.

**Table 1.**
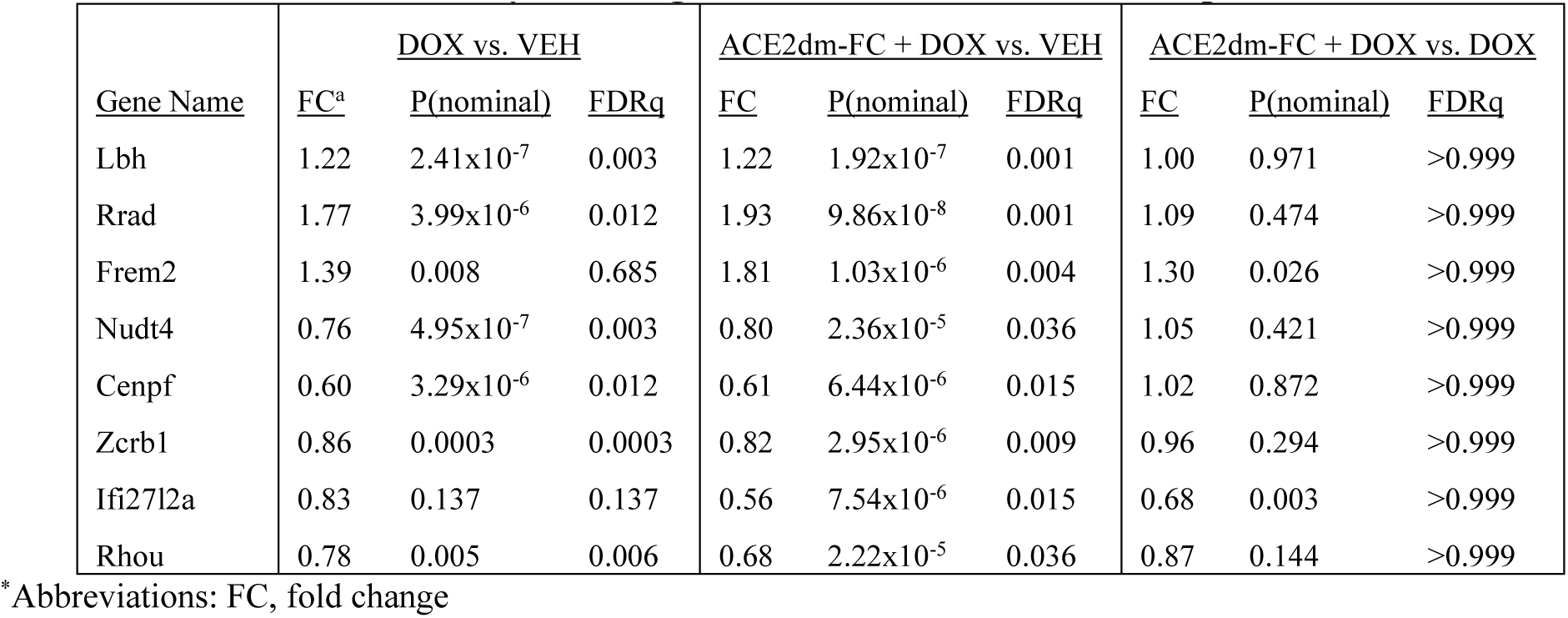
False Discovery Rate Significant Genes Between Comparisons.

|  | DOX vs. VEH |  |  | ACE2dm-FC + DOX vs. VEH |  |  | ACE2dm-FC + DOX vs. DOX |  |  |
|---|---|---|---|---|---|---|---|---|---|
| Gene Name | FC<sup>a</sup> | P(nominal) | FDRq | FC | P(nominal) | FDRq | FC | P(nominal) | FDRq |
| Lbh | 1.22 | 2.41x10<sup>-7</sup> | 0.003 | 1.22 | 1.92x10<sup>-7</sup> | 0.001 | 1.00 | 0.971 | >0.999 |
| Rrad | 1.77 | 3.99x10<sup>-6</sup> | 0.012 | 1.93 | 9.86x10<sup>-8</sup> | 0.001 | 1.09 | 0.474 | >0.999 |
| Frem2 | 1.39 | 0.008 | 0.685 | 1.81 | 1.03x10<sup>-6</sup> | 0.004 | 1.30 | 0.026 | >0.999 |
| Nudt4 | 0.76 | 4.95x10<sup>-7</sup> | 0.003 | 0.80 | 2.36x10<sup>-5</sup> | 0.036 | 1.05 | 0.421 | >0.999 |
| Cenpf | 0.60 | 3.29x10<sup>-6</sup> | 0.012 | 0.61 | 6.44x10<sup>-6</sup> | 0.015 | 1.02 | 0.872 | >0.999 |
| Zcrb1 | 0.86 | 0.0003 | 0.0003 | 0.82 | 2.95x10<sup>-6</sup> | 0.009 | 0.96 | 0.294 | >0.999 |
| Ifi27l2a | 0.83 | 0.137 | 0.137 | 0.56 | 7.54x10<sup>-6</sup> | 0.015 | 0.68 | 0.003 | >0.999 |
| Rhou | 0.78 | 0.005 | 0.006 | 0.68 | 2.22x10<sup>-5</sup> | 0.036 | 0.87 | 0.144 | >0.999 |

<sup>\*</sup>Abbreviations: FC, fold change

Utilizing differential gene expression data with more lenient cutoffs (|ΔFC = 1.5|, nominal P < 0.05), we identified the following differences with group-level comparisons: DOX vs. VEH, 36 upregulated and 42 downregulated; ACE2dm-Fc + DOX vs. VEH, 31 upregulated and 54 downregulated; ACE2dm-Fc + DOX vs. DOX, 29 upregulated and 27 downregulated (**Fig. 5A**). With this list from group-level comparisons, we applied Boolean logic to conduct overlap analysis and assign genes to 3 groups: common in doxorubicin treatment (*DOX Common*), treatment specific (*ACE2dm-Fc Specific*), and treatment responsive (**Fig. 5Bi**, supplemental data). We identified 21, 9, and 10 genes in the DOX common, treatment specific, and treatment responsive categories, respectively.

**Fig. 5.**
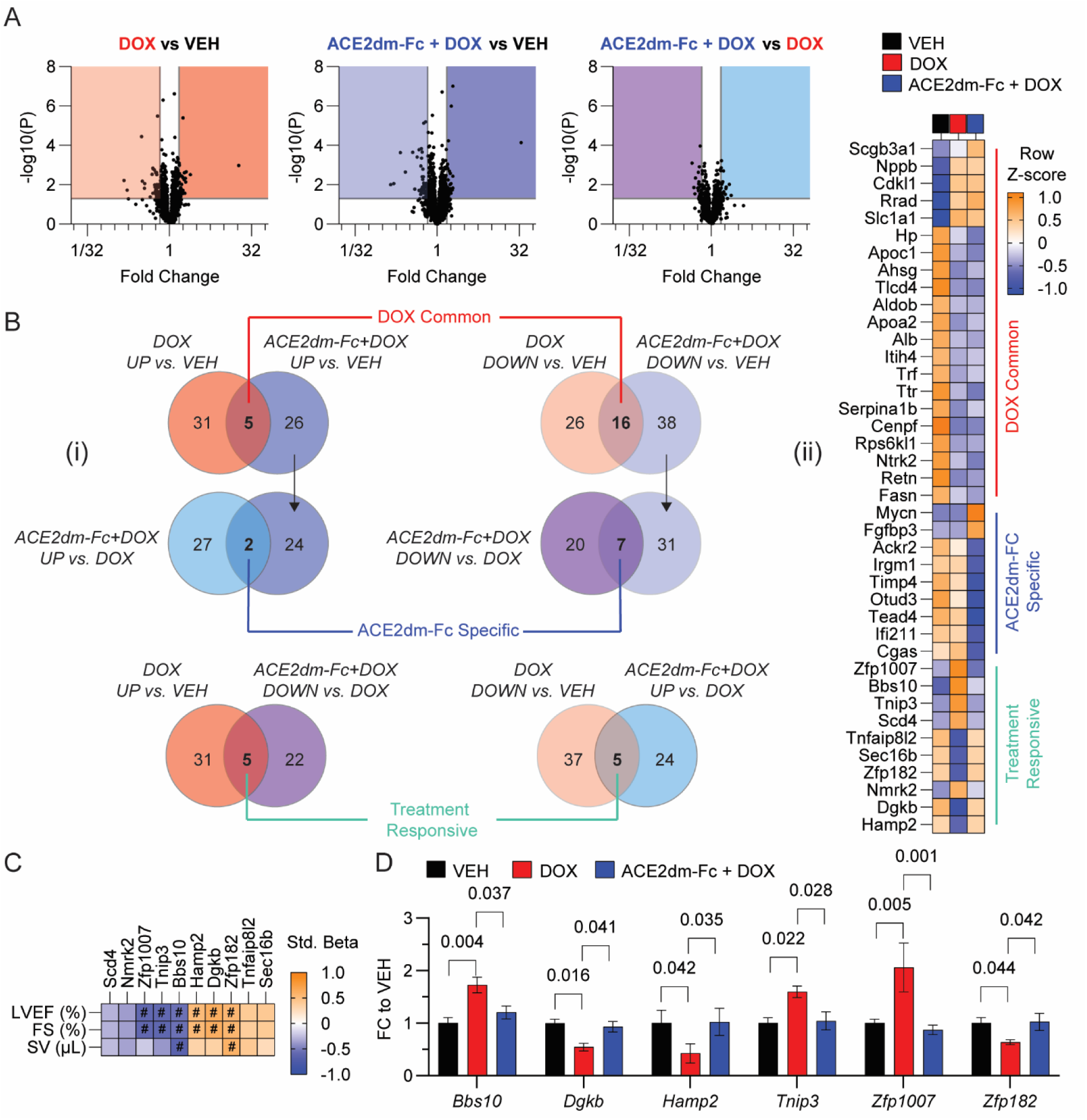
ACE2dm-Fc suppresses changes in transcription of genes that correlate with systolic heart dysfunction. (**A**) Volcano plots for each pairwise comparison. (**B**) Venn-Diagrams depicting Boolean/overlap analysis and heatmap of the resulting groups. (**C**) Correlations of gene expression and systolic function among treatment responsive genes and (**D**) those with FDR-level significance shown as FC to VEH. For all genes (**B-D**), |ΔFC = 1.5| and nominal p < 0.05. # indicates FDR < 0.05 correlation (row adjusted).

Finally, we identified gene expression differences between DOX and ACE2dm-Fc + DOX that correlated with multiple systolic function parameters. We reasoned that genes belonging to the treatment responsive set could represent correlates of mouse cardiac function independent of doxorubicin or ACE2dm-Fc administration (**Fig. 5B**). The genes included: *tnfaip8l2*, *sec16b*, *dgkb*, *hamp2*, *zfp182*, *nmrk2*, *bbs10*, *zfp1007*, *tnip3*, and *scd4* (**Fig. 5Bii**, supplemental data file). We found that 4 of these correlated with left-ventricular ejection fraction and fractional shortening (positive correlations: *amp2* and *dgkb*; negative correlations: *zfp1007* and *tnip3*) and 2 correlated with left-ventricular ejection fraction, fractional shortening, and stroke volume (positive correlation: *zfp182*; negative correlation: *bbs10*) (**Fig. 5C-D**, supplemental data file).

## DISCUSSION

There is a critical need for the identification of novel therapies that protect against cardiac dysfunction induced by anthracyclines such as doxorubicin. Standard-of-care heart failure medications targeting the RAS have demonstrated some benefit, but nuances regarding efficacy among patients suggest that further refinement could allow for broader cardioprotection among relevant clinical populations.^27,47^ Here, we provide evidence for a mechanistically distinct RAS modifying therapy, demonstrating that the novel engineered protein, ACE2dm-Fc is efficacious in a female mouse model of doxorubicin-induced cardiotoxicity by preserving LVEF, fractional shortening, stroke volume and reducing fibrosis.

Female mice were used due to the emphasis on translating ACE2dm-Fc to the clinic for use as an adjuvant in doxorubicin breast cancer chemotherapy regimens, particularly as women with breast cancer are ∼2-fold more likely to develop heart failure^14^ and because in the literature, mouse models of doxorubicin-induced cardiotoxicity have demonstrated sex differences.^33,39–41,48^ In our study, the functional cardioprotection imparted by ACE2dm-Fc treatment in systolic function aligns well with previous in vivo models of ACE inhibition in anthracycline-induced cardiotoxicity.^49–51^ ACE2dm-Fc treatment reduced animal heart rate under similar anesthetic exposure conditions to other groups (Fig. S1), which we suspect is related to vasodilative effects of MasR signaling.^52^ Lower animal heart rates can reduce cardiac function^53^ (e.g., especially in a purely mathematical sense for measures of cardiac output, which is definitionally unit volume-by-time), both mice treated with doxorubicin alone and doxorubicin + ACE2dm-Fc had lower cardiac output, with no difference among ACE2dm-Fc treated mice in cardiac output (Fig. S1). Given that mice treated with doxorubicin and ACE2dm-Fc had preserved stroke volume (the other contributor to cardiac output besides heart rate), we believe that potentially increased vasodilation from MasR signaling is not a cause for concern. Hypotension and drug intolerance constrain use of RAS-directed therapies in cancer patients. In the PROACT trial, which tested enalapril during high-dose anthracycline chemotherapy for breast cancer and lymphoma, 16% of patients discontinued the drug before completing chemotherapy, with symptomatic hypotension among the reasons, and four of these patients discontinued enalapril and chemotherapy concurrently.^22^ Entresto (sacubitril/valsartan) compounds this liability, pairing neprilysin inhibition, which raises circulating natriuretic peptides, bradykinin and adrenomedullin, with AT1R blockade by valsartan. This dual vasodilatory action produced more symptomatic hypotension than ACE inhibition alone in PARADIGM-HF (14.0% versus 9.2%) and in anthracycline-treated patients relative to placebo (14% versus 1.8%).^54^ Patients undergoing chemotherapy are especially susceptible given volume depletion from emesis and reduced oral intake, anemia, and concurrent antihypertensive therapy. ACE2dm-Fc has a different mode of action, enzymatically depleting Ang-II and generating Ang-(1-7) rather than blocking receptor signaling, and the preservation of stroke volume without further reduction in cardiac output relative to doxorubicin alone argues against clinically meaningful hemodynamic compromise.

Su *et. al.* found that female mice had fewer differentially expressed genes compared to males following treatment with doxorubicin. Compared to our study, they found a higher number of DEGs but used a more acute endpoint and a different mouse background.^55^ With FDR adjustment, we identified 4 gene expression changes common to both doxorubicin treated groups. Among those, *cenpf*, a DNA damage response gene, has previously been identified to greatly decrease in iPSC cardiomyocytes treated with doxorubicin at acute and chronic treatment timepoints (2-12 days),^56^ consistent with our observation of transcript depletion.

As FDR correction yielded no significant differences in transcriptomic programming between DOX and ACE2dm-Fc + DOX, we used two orthogonal approaches to identify correlates of cardiac function that could reveal subtle differences between the two groups. Since we could not perform more stringent thresholding at the individual transcript level, we felt it was prudent to perform statistical test correction for multiple regression analyses. Discriminant analysis (i.e. sPLS-DA) resulted in the identification of known effectors of cardiac pathology during doxorubicin treatment, *nr4a1* and *btg2*. Nur77, the protein product of *nr4a1*, is an orphan nuclear receptor that plays roles in cardiac remodeling via NLRP3 inflammasome activation. In vivo studies of doxorubicin-induced cardiotoxicity demonstrated that *nr4a1* promotes cardiac dysfunction, fibrosis, and cardiomyocyte injury, which were ablated following knockdown.^57^ Similarly, we saw that increased *nr4a1* expression negatively correlates with systolic function. Ang-II can acutely induce transient upregulation of *nr4a1*/Nur77, but these investigations used stressors other than doxorubicin, indicating that *nr4a1*’s role in cardiac protection or injury could be stress and context dependent.^58–60^ The relationship with *btg2* is clearer, as several investigations have shown it is upregulated in the heart in response to doxorubicin and that reducing total protein levels (with drugs or micro-RNAs) reduces cardiotoxicity and improves heart function.^61–63^ *btg2* has previously been implicated as a potential therapeutic target for chemotherapy induced cardiotoxicity.^64^

Additional investigation of the treatment responsive genes identified in this study, their roles in cardiac pathology and how they may be altered by RAS modulation, could contribute to better understanding of how they could be targeted for heart dysfunction. We further posit that ACE2dm-Fc reduces myocardial energetic and pathophysiological stress in the context of doxorubicin such that the activation of the stress response genes, *nmrk2* and *tnip3*, are not required. This is consistent with transcript levels in vehicle-treated mice, where no cardiac stress was introduced. *nmrk2* and *tnip3* have known associations with heart failure; however, *mmrk2* did not correlate with mouse heart function. Nmrk2 is an NAD+ salvage enzyme critical for protection against dilated cardiomyopathy.^65^ Preservation of NAD+ (and related metabolites) pools via the salvage pathway serves a cardioprotective role following doxorubicin.^66^ Tnip3 has not been previously implicated in doxorubicin induced cardiac injury but has an established link to cardiac health. Shi *et. al.* saw upregulation of *tnip3* following transverse aortic constriction, but knock-out lead to greater pathological remodeling. They went on to show that Tnip3 plays an important role in reducing cardiomyocyte hypertrophy via STAT1 stabilization and reduced NF-Κβ activation.^67^ Bbs10, which we observed to negatively correlate with systolic function, has transient links to cardiovascular disorders and RAS signaling. Related genes are mutated in Bardet-Biedel Sydrome (BBS), an autosomal recessive inheritance patterned pleiotropic genetic condition where patients experience renal abnormalities, hypertension, and obesity, which are risk factors for cardiovascular disease and related to RAS.^68,69^ Other correlates of systolic function we identified have less clear relevance to cardiovascular dysfunction.

Study limitations include the absence of a mouse group treated with ACE2dm-Fc alone, which precluded us from isolating the independent effects of ACE2dm-Fc. It is possible that ACE2dm-Fc enhances basal LV systolic function, thus providing generalized cardiac functional enhancement as opposed to resilience against doxorubicin injury. An additional consideration is that transcripts identified in discriminant and overlap analyses to identify gene groups were not FDR adjusted. Also noteworthy is that ideally, novel cardioprotective treatment regimens cause no change in tumor size or chemo-sensitizing effects while also preserving cardiac function. As we did not utilize a tumor bearing model, we could not evaluate ACE2dm-Fc regarding tumor changes. Preclinical studies so far suggest that RAS modulation attenuates tumor growth.^70–72^

Despite these limitations, our findings demonstrate great potential for this novel approach of RAS modulation to serve as a cardioprotective therapy for doxorubicin-induced cardiotoxicity. In future work, alternative dosing routes, such as subcutaneous and intravenous delivery, and their pharmacodynamics should be established. Multiple experimental endpoints, especially earlier following ACE2dm-Fc administration, to establish a shift towards MasR directed signaling in relevant tissue types (as these may be expected to occur in lung, liver, and kidneys) that could promote circulatory or systemic effects, should be utilized to more cleanly delineate the proposed mechanism of action.

## Acknowledgments

We thank Brandy Edenfield and Jenni Pattineik of the Mayo Clinic Histology Cores for preparing tissue blocks and slides.

## Author contributions

Conceptualization: NN, PH, NKHN, DN

Methodology: NN, PH, NKHN, DND, MRF, CRW

Investigation: NKHN, DND, CRW, IEP, OAS, MRF

Visualization: DND, PH

Funding acquisition: NN, PAR

Project administration: NN, PAR, MF, DND

Supervision: NN, PH, MRF, PPA, CPR, DND, NKHN

Writing - original draft: DND, NKHN, NN, MRF, PH

Writing - review & editing: DND, NKHN, CRW, IEP, OAS, CPR, PPA, MRF, PH, NN

